# The Culicinae are Monophyletic and Ancient: A response to Pierce et al. 2025

**DOI:** 10.64898/2026.05.04.720205

**Authors:** John Soghigian, Gen Morinaga, Huiqing Yeo, Yvonne-Marie Linton, Richard C. Wilkerson, Maria Anice Sallum, Igor Sharakov, Maria Sharakova, Magdalena Laurito, Woo Jun Bang, Seunggwan Shin, Loki Snyman, Thomas Zavortink, Charles Sither, Michael Reiskind, Brian M. Wiegmann

## Abstract

Mosquitoes are classified into two subfamilies, each monophyletic, and typically considered to both be ancient, having diverged more than 100 million years ago based on previous divergence analyses. A recent publication challenged this view with phylogenomic results primarily from the third codon position and UCEs. Utilizing alternative fossil placement and these phylogenomic data, these authors find that the Culicidae and Chaoboridae diverged in the lower Cretaceous, and that one mosquito subfamily, the Anophelinae, is nested within the Culicinae. These results are in stark contrast to previous results from diverse data sources, ranging from other genomic data, to morphology, to fossils. Here, we briefly detail the substantial evidence that supports two monophyletic subfamilies of extant mosquitoes, along with fossil evidence that supports the ancient divergence of these lineages.

## Main Text

Two monophyletic subfamilies of extant mosquitoes are currently recognized: the Anophelinae, with approximately 500 species in 3 genera, and the Culicinae, with approximately 3,200 species in 11 tribes and 39 genera(1, 2). Recently, Pierce et al.(3) (PEA) challenged this view with phylogenomic results derived primarily from the third codon position (NT3) and UCEs showing the Anophelinae nested within the Culicinae, with the subfamily Anophelinae and the genus *Culex* sister to one another (“ACS”, in contrast with a two subfamilies hypothesis, “TSH”). PEA also found that Culicidae and Chaoboridae diverged in the lower Cretaceous, far younger than previous estimates. We are unconvinced by these results and find instead that the overwhelming body of evidence strongly supports an ancient and monophyletic Culicinae.

There is widespread support from diverse sources for the TSH. Analyses on morphological data have consistently recovered a monophyletic Culicinae, based on eight synapomorphies(2), while there are no synapomorphies found between *Culex* and the Anophelinae alone(2) (Fig.1). Non-coding markers such as 18S(4), and mitochondrial genomes(5), both support two monophyletic subfamilies. Gene families, such as odorant binding proteins(6), and their respective copy numbers(6), also reflect a monophyletic Culicinae. Previous phylogenomic analyses also have supported the TSH(7).

**Figure 1.**
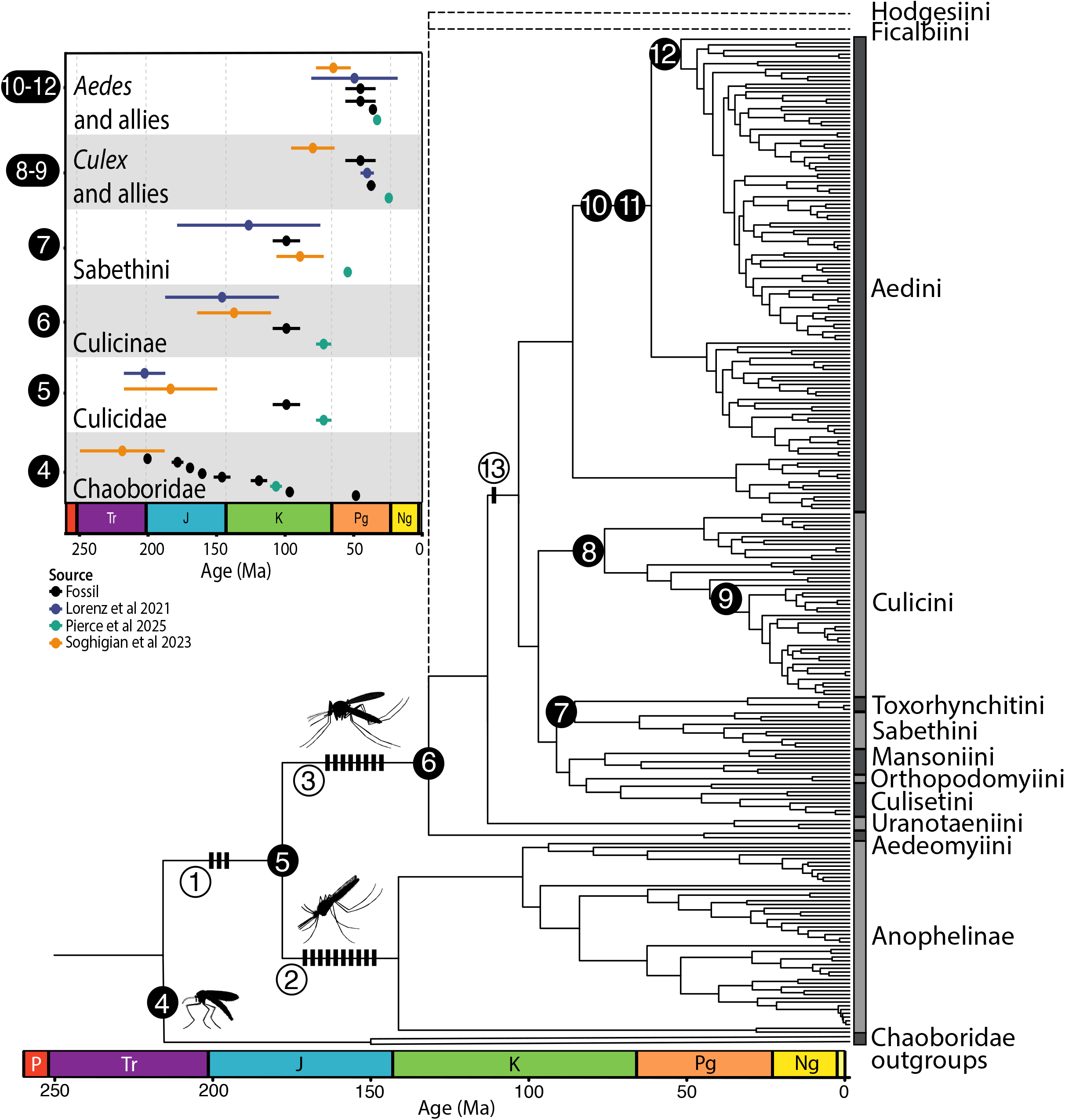
The phylogeny of the Culicidae based on previous publications, with unsampled tribes shown as dotted lines (2, 7). Open circles with numbers and thatched lines on branches indicate synapomorphies identified for the Culicidae (1 – 3 synapomorphies), the Anophelinae (2 – 10 synapomorphies), and the Culicinae (3 – 8 synapomorphies)(2). The inset compares fossil ages(1, 9, 13) with divergence times from three studies(3, 5, 7). Dark colored numbers correspond to approximate fossil placement on extant lineages, except for the Chaoboridae, which is based on the age of the divergence between mosquitoes and Chaoboridae, as many Chaoboridae fossils are in extinct or unsampled lineages, and for the Sabethini, which was estimated based on the common ancestor of the Sabethini and its sister lineage, due to sampling differences between relevant studies. *Aedes* and allies here refers to aedine genera other than *Psorophora*(7).

PEA argues that branch attraction bias creates the TSH, where fast-evolving Anophelinae draw long branch outgroups towards them. However, we believe ACS topology is a result of compositional heterogeneity at NT3. In our previously published analyses (7), outgroups and some mosquitoes have low-GC at NT3 but not other positions (Fig.2A), as also observed in PEA. Accounting for GC heterogeneity, such as with using NT1/2 or amino acids (Fig.1), recoding NT3 as a purine or pyrimidine(8) (Fig.2C), or using higher-GC outgroups but the same ingroups (8) (Fig.2D), recovers two monophyletic subfamilies. This is consistent with branch attraction owed to GC content heterogeneity at NT3 causing the ACS topology.

**Figure 2.**
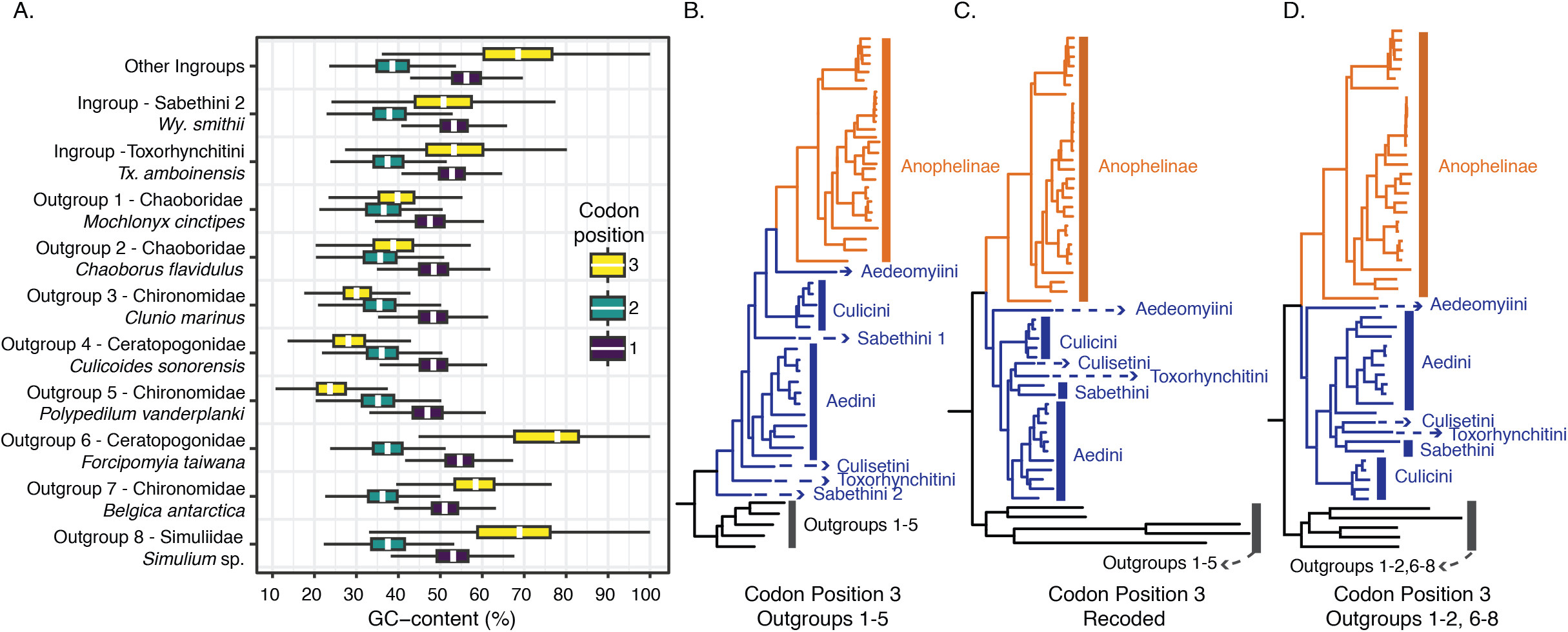
A) A boxplot of GC content from an alignment of 5,667 genes from 55 genomes and transcriptomes (7, 8). Outgroups 1-5 were used in Soghigian et al. 2023 (7), while outgroups 6-8 are higher GC content outgroups used to illustrate the effects of GC-content heterogeneity. B) The phylogenetic analysis of NT3, following methods described in Soghigian et al. 2023 (7) from a previously published dataset of 5,667 genes from 55 genomes and transcriptomes (7). C) The phylogenetic analysis of the same NT3 dataset as in B, but with purines recoded as R and pyrimidines recoded as Y. D) The phylogenetic analysis using the same ingroup species as in B, but with outgroups (replacing outgroups 3-5 with 6-8, see panel A) whose GC content is more similar to mosquitoes. All branch support values between tribes and mosquito subfamilies are 100.

We also believe that the divergence times presented by PEA are too young, likely owed to differences in priors from past studies (e.g., distributions and fossil placement). Crown fossils placed in extant mosquito lineages(1) are significantly older than the divergence times reported for those genera in PEA(Fig.1B). The ages of fossils such as *Culex* (*Culex*) *erikae*(1), *Aedes* (*Ochlerotatus*) *serafini*(1), and *Cretosabethes primaevus*(9), are more consistent with previous divergence time analyses(Fig.1B) than with PEA. Notably, the latter fossil is older (∼99 MYA) than PEA’s estimate for the common ancestor of all extant mosquitoes.

Further, we caution that >85% of Diptera UCEs(10), such as those used by PEA, are in protein coding regions. Failure to properly partition these genic markers, particularly with flanking non-genic sequences included, could result in model misspecification and mislead both phylogenetic and dating analyses.

In conclusion, we remain unconvinced by the results of PEA. Compositional heterogeneity among fast-evolving sites like NT3s can lead to long-branch attraction and thus incorrect topologies, which explains the ACS. We believe the overwhelming body of evidence currently supports two extant subfamilies of mosquitoes, each ancient and monophyletic, and that this support comes from a range of sources—from morphology to fossils to genomes.

## Methods

We retrieved morphological characters from a previous publication (2) and compared whether these characters supported either phylogenetic hypothesis of mosquito evolution: the two-subfamily hypothesis (TSH) or the *Anopheles*-*Culex* Sister hypothesis (ACS). Then, we placed relevant synapomorphies on the phylogenetic tree of the Culicidae from Soghigian et al. 2023 (7). We retrieved fossil ages from relevant repositories, and compared their ages to clade ages from three publications: Pierce et al. (3), Soghigian et al. (7), and Lorenz et al (5).

To evaluate nucleotide content heterogeneity, we used previously published and/or publicly available data, as the alignments analyzed by Pierce et al. (3) were not publicly available. GC content per codon position was recorded from 5,667 genes from 55 genomes and transcriptomes previously published (7), then visualized as a barplot in R v 4.3 with ggplot (11). We reconstructed phylogenies from nucleotide codon position 3 alone using IQ-Tree 2(12), as this had not been done with this previously published dataset. Then, we recoded the third codon position following a so-called RY recoding strategy, wherein purines are recoded as R and pyrimidines as Y, and re-analyzed the alignment with IQ-Tree again. As some outgroup species had particularly low GC content, an additional phylogenetic analysis was performed in which these low GC content outgroups (*Clunio marinus, Culicoides sonorensis*, and *Polypedilum vanderplanki*) were removed and replaced with other Culicimorph outgroups with higher GC content. After retrieving proteomes for these species from public repositories, we followed previously published methods (7) to align these new outgroups with the remaining outgroups and ingroups. We then performed phylogenetic analysis on these new alignments, which had the same ingroup as all previous analyses, but three different outgroups, with IQ-Tree.

## Data Availability Statement

Alignments and associated data used here was previously published (7) and/or are publicly available (*Simulium* sp - GGBP00000000, *Forcipomyia taiwana* - GCA_963930915.1, *Belgica antarctica* - GCA_000775305.1). Code, data, and other associated files are available on https://github.com/jsoghigian/culicitree/tree/main/comphet/ and https://github.com/jsoghigian/culicitree/tree/main/comphet/Soghigianetal2026.

## Acknowledgements

We thank G. Powell, R. Lanfear, J. Steenwyk, J. Gilleard, H. Schaezl, and the Sperling Lab at University of Alberta for helpful discussions related to this letter and/or comments on previous drafts. We also thank the Research Computing Services group at the University of Calgary for access to the ARC cluster, where analyses were performed. We acknowledge the support of the Natural Sciences and Engineering Research Council of Canada (NSERC), funding reference number RMS21-73779779 [Cette recherche a été financée par le Conseil de recherches en sciences naturelles et en génie du Canada (CRSNG), numéro de référence RMS21-73779779].

